# Evolution of Tn*4371* family ICE; *traR* mediated coordination of cargo gene upregulation and horizontal transfer

**DOI:** 10.1101/2024.03.05.583556

**Authors:** Satoshi Matsumoto, Kouhei Kishida, Shouta Nonoyama, Keiichiro Sakai, Masataka Tsuda, Yuji Nagata, Yoshiyuki Ohtsubo

## Abstract

ICE_KKS102_Tn*4677*, which has been shown to transfer horizontally, carries *bph* operon for mineralization of PCBs/biphenyl and belongs to an ICE Tn*4371* family. In this study we investigated the role of *traR* gene encoding a LysR-type transcriptional regulator, which is conserved in sequence, positioning, and directional orientation among Tn*4371* family ICEs. The *traR* belonged to *bph* operon and its overexpression on solid medium resulted in modest upregulation of *traG* (3-fold) and marked upregulation of *xis* (80-fold), and enhanced ICE excision, and notably ICE transfer frequency. We propose the evolutional roles of *traR*, which upon insertion to the current position, connected the cargo gene activation and ICE-transfer. This property of ICE, transferring under environmental conditions that lead to cargo gene activation, would give fitness advantages to the host bacteria, thereby resulting in efficient dissemination of the Tn*4371* family ICEs.

**Significance:** Only ICE_KKS102_Tn*4677* is proven to transfer among the widely disseminating Tn*4371* family ICEs from β and γ-proteobacteria. We showed that the *traR* gene in ICE_KKS102_Tn*4677* conserved in the ICE family with fixed location and direction is co-transcribed with the cargo gene and activates ICE transfer. We propose that capturing of *traR* by an ancestral ICE to the current position established ICE Tn*4371* family ICEs. Our findings provide insights into the evolutionary processes that led to the widespread distribution of the Tn*4371* family of ICEs across bacterial species.

## INTRODUCTION

Mobile genetic elements play a crucial role in bacteria in acquiring new traits, including antibiotic resistance, symbiosis, virulence, and compound degradation (1) (2) (3) (4). Integrative and conjugative elements (ICEs) are among the various types of mobile genetic elements, which are sometimes referred as plasmids integrated in the bacterial chromosome. ICEs excise themselves from the chromosome, form a plasmid like circular structure, transfer to a recipient cell by conjugation, and reintegrate into the recipient chromosome (5).

Followings are the molecular mechanisms of ICE conjugative transfer with the RP4 plasmid system nomenclature where applicable(6). Integrase (encoded by *int* gene) with the aid of excisionase (encoded by *xis* gene) mediates excision by catalyzing site-specific recombination between *attL* and *attR*, which locate at left and right boundary of ICE. A set of two sites *attP* and *attB* are generated by the recombination, *attP* is on a plasmid-like circular entity and *attB* is on a chromosome. The plasmid form of ICE is transferred to recipient cell by Dtr and MPF (<u>m</u>ating-<u>p</u>air formation) systems, which are extensively characterized in number of studies on plasmid transfer. In the transfer, *oriT* (origin of transfer) is recognized and nicked by relaxase (encoded by *traI* gene) to form relaxosome, in which relaxase is covalently bound to 5’ end of DNA. The DNA-relaxase complex is passed to MPF system by the function of the coupling protein (encoded by *traG*). The MPF system, resembling to type IV secretion system, exports the complex of relaxase and single-stranded DNA, which is released as the progression of rolling-cycle replication, into a recipient. In the recipient cell the imported DNA is recircularized and revert back to double strand DNA, and then integrated into the chromosome by the integrase.

Among diverse ICE families is Tn*4371* family of ICEs. Tn*4371* is 54.7-kb transposon carrying PCB/biphenyl degrative genes (*bph* genes) and isolated after mating assay from *Cupriavidus oxalaticus* A5 carrying a broad host range plasmid RP4 and *Cupriavidus matallidurans* CH34 (7). In a 2003 study by Ariane Toussaint et al., the biphenyl catabolic transposon Tn*4371* was thoroughly sequenced and analyzed, revealing its mosaic structure with multiple building blocks, and a new family of genomic island was proposed(8). Although Tn*4371* has never been proven to transfer conjugally, the designation of ICE Tn*4371* family has been used to include a growing number of ICEs (9) (10) (11) (12). Ohtsubo et al. compared 112 related elements and showed that βγ-type ICEs (a sub group of Tn*4371*-related ICEs from β and γ-proteobacterial species) have distinct conserved structure and distinguishable from those from α proteobacteria (αI and αII types). The βγ-type ICEs (Tn*4371* family ICEs) carry four modules include *int, parB*, *traI* and *mpf* blocks (13).

Tn*4371* family ICEs share the same modular structure, and notably a *traR* gene, encoding LysR family transcriptional regulator, is conserved as the first gene of the *mpf* block, which encodes type four secretion system (T4SS) components (13, 14). This positioning, coupled with its sequence conservation among Tn*4371* family ICEs, suggests its important role in the dissemination of Tn*4371* family ICEs.

The ICE_KKS102_Tn*4677* (Fig.1) identified from PCB/biphenyl degradative bacterium *Acidovorax* sp. KKS102(15) is the only ICE in the Tn*4371* family for which horizontal transfer was confirmed (13). The ICE is 61.8 kb in length and located upstream of the Gly-tRNA gene and flanked by 9 bp *attL* and *attR* direct repeats (GATTTTAAG). ICE_KKS102_Tn*4677* carries *bph* genes for degradation of PCB/biphenyl. The expression of the *bph* genes is dependent on p*E* promoter, whose activity is repressed by BphS through its binding to operator sites overlapping the transcription start site of *pE* (16). The repression by BphS is alleviated by 2-hydroxy-6-oxo-6-phenylhexa-2,4-dienoic acid (HOPDA), a metabolite of the biphenyl degradation pathway (17) (16). The activity of *pE* promote is also regulated by favorable organic compounds, for which a two-component regulatory system composed of BphP and BphQ is involved (18) (19).

**Fig. 1.**
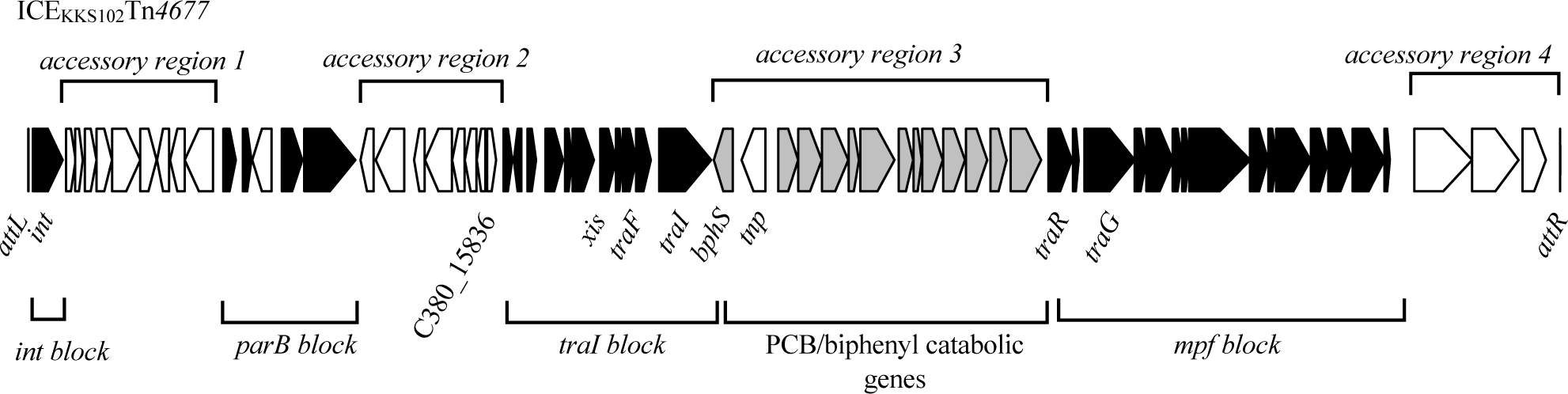
Structure of ICE_KKS102_Tn*4677*. The black arrows represent conserved genes, grey arrows represent PCB/biphenyl catabolic genes, and white arrows represent accessory genes.

The transfer of ICE_KKS102_Tn*4677* to α, β, and γ proteobacterial strains was confirmed, but it occurred at a very low frequency (about 10^-10^ per donor). This low transfer frequency is partly attributed to the low proportion of the ICE being in its excised form (one excised form among 10^5^ cells) (13).

In this study, we investigated the role of *traR* gene in conjugation. We demonstrated that *traR* gene is transcribed from the *pE* promoter, and that *traR* gene product activates the transfer of ICE_KKS102_Tn*4677* by increasing the excised form by activating the transcription of *xis*. The excision alone, however, did not explain the activation of transfer as *xis* overexpression did not activate the transfer, and as the transfer of an *oriT* plasmid required *traR* overexpression. Identification of multiple possible TraR binding sites suggested pleiotropic activation function of TraR in conjugation. We discuss the evolutional role of *traR* in the dissemination of Tn*4371* family ICEs.

## MATERIALS AND METHODS

### Strains and growth condition

The bacterial strains and the plasmids used in this study were listed in Table 1. *Escherichia coli* strains were grown at 37<C in LB broth and the other strains were grown at 30<C in 1/3 LB broth. Cells carrying the plasmid were cultured in a medium supplemented with antibiotics; 100 µg/mL for ampicillin, 50 µg/mL for Kanamycin (Km), 10 µg/mL for Gentamicin (Gm), and 5 µg/mL for Tetracycline (Tc).

**Table 1.**
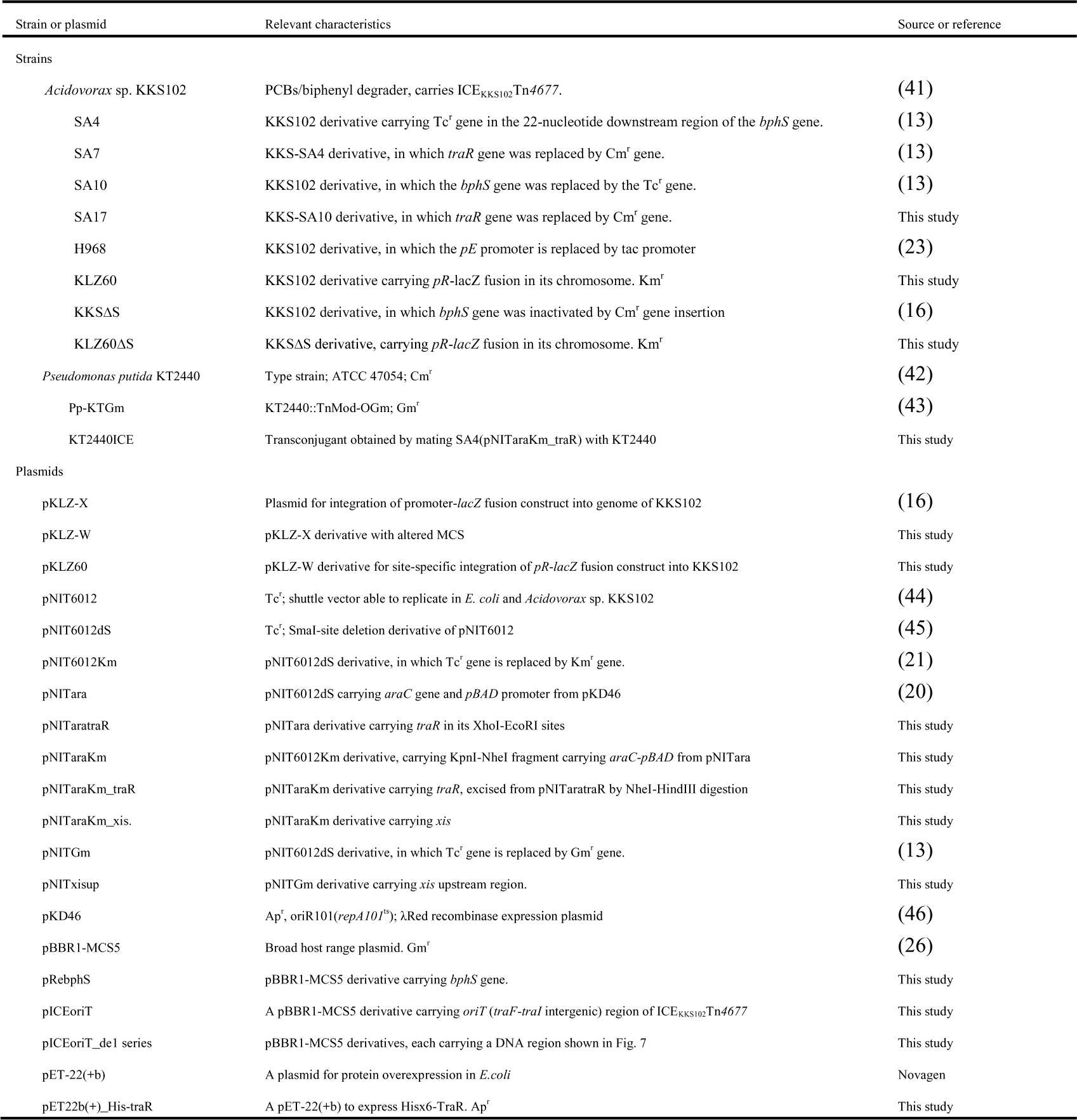
Strains and Plasmids.

### Primers

The primers used in this study are listed in Table S1.

### Construction of plasmids and strains

For the construction of KLZ60, a plasmid pKLZ60 was constructed by amplifying 1 kb *traR* upstream region by using a pair of primers KKS-pR-F and KKS-pR-R, and ligation of it to *lacZ* gene on a reporter integration plasmid, pKLZ-W. The pKLZ-W was created by digestion of pKLZ-X(16) with EcoRI and BamHI and subsequent ligation of a DNA fragment prepared by annealing a pair of synthetic DNAs, PKLZ-W-a and PKLZ-W-b. The pKLZ60 was used after linearization by HindIII digestion to obtain strains, in which the pR promoter-*lacZ* construct was inserted into a specific location in the chromosome by homologous recombination. We obtained KLZ60 and KLZ60ΔS, by introducing linearized pKLZ60 into KKS102 or *bphS* disruptant(16), respectively. The plasmid pRebphS was created by cloning of *bphS* containing fragment into BamHI and HindII sites of pBBR1-MCS5.

The plasmid pNITaraKm was created by transferring *araC-pBAD* fragment of pNITara (20) into pNIT6012Km (21). The plasmid pNITaratraR was constructed by inserting a DNA fragment amplified by using a pair of primers traR-GA-F and traR-GA-R into pNITara(20). The cloned fragment of pNITaratraR containing *traR* was excised by NheI-HindIII digestion and ligated into pNITaraKm to create pNITaraKm_traR. The plasmid pNITaraKm_xis was constructed by cloning of a DNA fragment amplified by a pair of primers SM083 and SM084 into pNITaraKm. The pNITxisup was constructed by cloning of a DNA fragment amplified by a pair of primers SM103 and SM104 into pNITGm.

A series of pICEoriT plasmids were constructed by cloning PCR-amplified fragments into EcoRI-HindIII sites of pBBR1-MCS5. The cloned regions are depicted in Fig. 7.

The plasmid pET22b(+)_His-traR for the expression of N-terminal Hisx6-tagged TraR was constructed by amplifying *traR* gene fragment with a pair of primers SM141 and SM142, and cloning of it into NdeI-XhoI site of expression plasmid pET-22b(+) (Novagen, WI, USA).

### Mating experiments

For mating assay, donor and recipient cells were grown in 5 mL of 1/3 LB medium for 18 hours. Subsequently, cells in 1 mL of each culture were harvested, washed once with fresh medium, mixed and suspended in 50 µL of fresh medium. This suspension was spotted onto solid 1/3 LB media. To induce *traR* gene expression cloned under *pBAD* promoter, solid 1/3 LB media containing 10 mM arabinose was used. After incubation at 30<C, cells were collected and plated onto solid media containing antibiotic(s) for selection.

### LacZ activity measurement

LacZ activity was measured as described previously (16).

### Quantification of circular form of ICE

After strains were grown up to the end of the log phase, cells in 1 mL of the culture were harvested by centrifugation and suspended in 50 µL of fresh medium. The suspension was spotted on a solid 1/3 LB medium containing 10 mM of arabinose to induce *traR* expression. After incubation, cells were resuspended in MiliQ water and incubated at 95℃ for 10 minutes. After cells were removed by centrifugation, supernatants were used as templates for quantitative Real-Time PCR (qRT-PCR).

qRT-PCR was conducted by using Luna Universal qPCR Master Mix (New Englad Biolabs) and a CFX Connect^TM^ Real-Time System (Bio-Rad Laboratories, Hercules, CA). The PCR conditions are initial 2 minutes at 95<C, 40 cycles of 10 seconds at 95 <C, 10 seconds at 61.4 <C, and 10 seconds at 68<C. A set of primers SM349 and SM350 was used for the detection of *attB*, SM025-SM026 for *attP*, and SM026-SM027 for the both of the ICE forms. For each sample, the three sets of primers were used. The results obtained with SM026-SM027 served as an internal standard to normalize the differences in the amount of DNA used for the qPCR.

### Quantification of the expression of genes in ICE_KKS102_Tn*4677*

RNA in each sample was extracted by ISOGEN^TM^ (NIPPON GENE) according to the provided protocol. After extraction, each RNA sample was treated with DNase I (TAKARA) at 37<C for 2 hours to remove the residual DNA. Reverse transcription was conducted by using ReverTra Ace®□ qPCR RT Master Mix (TOYOBO) and random 9-mer primers.

The qRT-PCR was conducted as described for the detection of *attP* and *attB*. Sets of primers SM015-SM016, SM019-SM020, SM041-SM042, SM105-SM106, and SM142-SM143 were used for detection of *traR*, *rrn*, *traG*, *xis*, and *traI*, respectively. The expression levels of each gene were normalized to the expression levels of the rRNA gene, *rrn*.

### Purfication of TraR

TALON Metal Affinity Resin (TAKARA, Shiga, Japan) was used to purify N-terminal Hisx6-tagged TraR protein, according to the provided protocol.

### Gel shift assay

The FAM-labeled *xis*-upstream-region DNA was prepared by a pair of primers PNIT5041-FAM and PNIT5548-FAM using pNITxisup as a template. The FAM-labeled *oriT*-region DNA was prepared by a pair of primers M4out-FAM and RVout-FAM using pICEoriT as a template. The FAM-labeled *traG*-upstream-region DNA was prepared by a pair of primers SM484 and SM485-FAM using genomic DNA of KKS102 as a template. For the preparation of DNA containing the TraR binding motif, the following pairs of oligonucleotides were mixed and annealed: X001-X002 (BSxis), X003-X004 (BStraG), X005-X006 (BStraI), and X007-X008 (BStraR).

## RESULTS

### Strains used in this study

Strains used in this study are listed in Table 1. Specifically, the SA4 strain is identical to the wild-type strain KKS102, except that it carries a tetracycline resistance gene inserted downstream of the *bphS* gene. The SA10 strain is identical to the SA4 strain, except that the *bphS* gene is replaced with the tetracycline resistance gene.

### *traR* belongs to the *bph* operon and its transcription is regulated by *bphS*

We predicted that *traR* is part of *bph* operon because it is located downstream of *bph* genes in the same orientation. To test this, we conducted qRT-PCR to measure the *traR* transcription level in a *bphS* deletion strain (strain SA10), and in a strain in which *pE* promoter for *bph* operon is replaced by a constitutive *ptac* promoter strain KH968 (Fig. 2, panel A) (22) (23). The qRT-PCR analysis revealed a 13-fold increase of *traR* transcripts in SA10, and this increase was suppressed by complementing SA10 with pRebphS, a plasmid carrying *bphS*. Similarly *traR* transcription was elevated 30-fold in KH968. These observations showed that *traR* is part of the *bph* operon and transcribed from *pE* promoter.

**Fig. 2.**
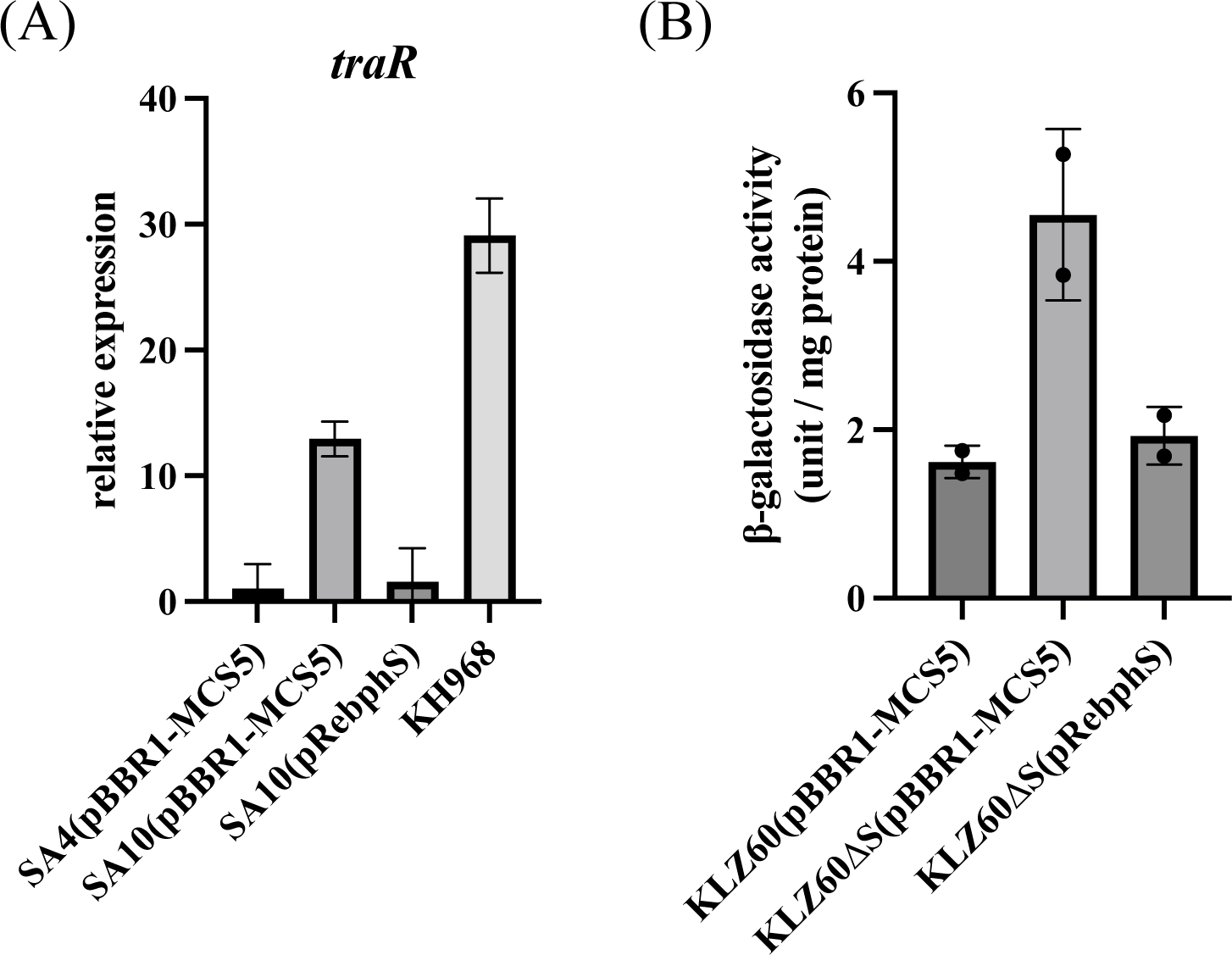
*traR* belongs to *bph* operon. (A) qRT-PCR assay showing that *traR* belongs to the *bph* operon. Each strain was grown to a OD600 value of 0.8 in 1/3 LB medium, and then total RNA was isolated for qRT-PCR analysis. The values are relative value to the strain SA4(pBBR1-MCS5), and *rrn* level was used as internal standard. (B) LacZ reporter assay showing that *pR* promoter level is increased in Δ*bphS.* LacZ activity is expressed as unit/mg protein (16). Each strain was grown as in panel A.

### *pR* promoter for *traR* transcription is upregulated in Δ*bphS*

To test the possibility that *traR* has its own promoter and to test if the promoter is regulated by *bphS*, 1 kb upstream region of *traR* was ligated in front of *lacZ* gene and integrated into the genome of KKS102 (resulting in a strain KLZ60) to measure the promoter activity of this region (Fig. 2, panel B). We found that the LacZ activity was increased when *bphS* gene in KLZ60 was deleted (strain KLZ60ΔS), and this increase was suppressed by supplying *bphS* gene in trans. These results indicated that a promoter is present in front of *traR* (designated as *pR* promoter), and that *pR* is negatively regulated by BphS.

### *traR* overexpression activates the transfer of ICE_KKS102_Tn*4677*

To test whether *traR* expression activates the transfer of ICE_KKS102_Tn*4677,* we transformed SA4 with pNITaraKm_traR. SA4 (pNITaraKm_traR) was mated with *P. putida* KT2440Gm (a KT2240 derivative carrying gentamycin resistance gene for selection) in the presence of different concentrations of arabinose. As shown in Fig. 3, a significant increase in the transfer frequency of ICE_KKS102_Tn*4677* was observed in an arabinose-concentration dependent manner, showing that the *traR* expression activates the ICE transfer. The colonies grown on a plate containing Tc and Gm were confirmed that they carry ICE_KKS102_Tn*4677* by colony PCR (data not shown). No transfer was observed in the vector control experiment using SA4 (pNITaraKm).

**Fig. 3.**
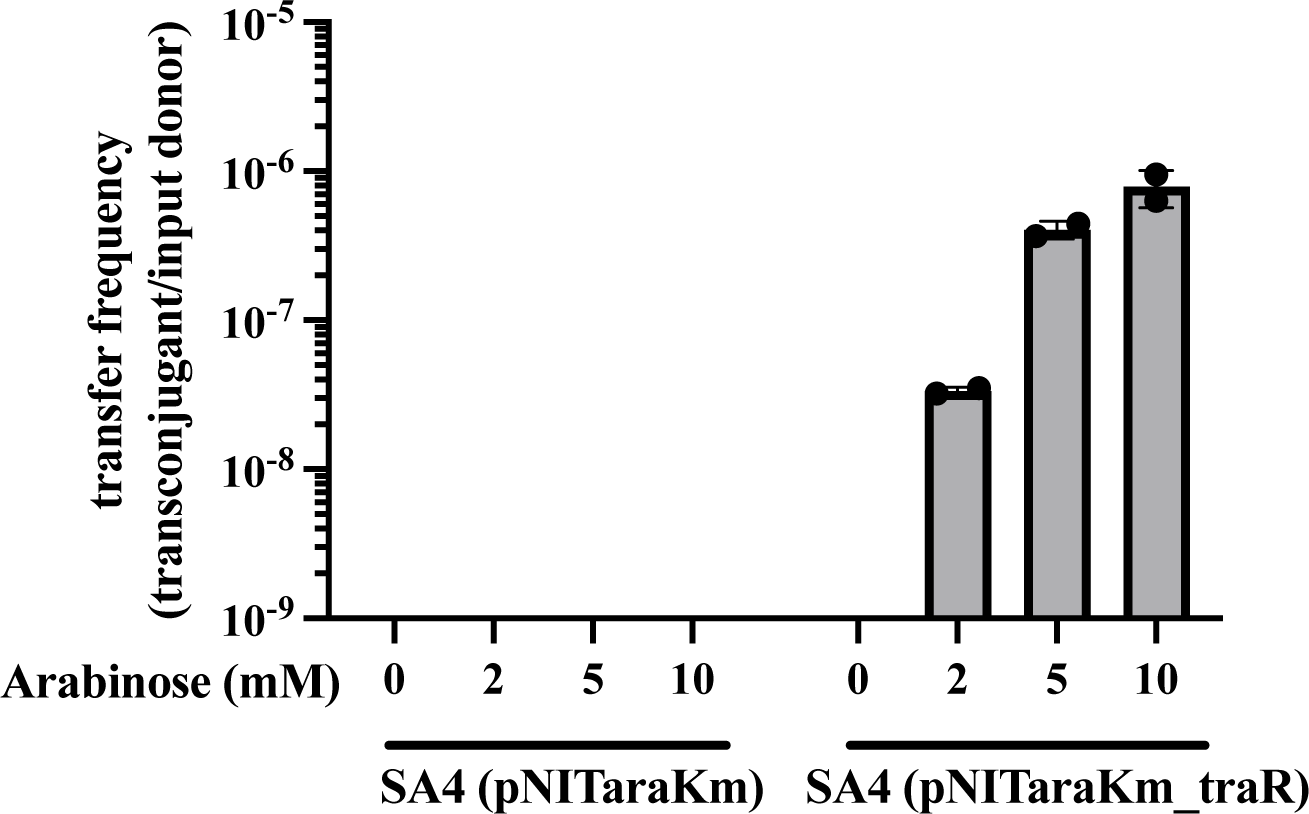
Overexpression of *traR* increased ICE transfer frequency. Donor strains [SA4 (pNITaraKm_traR) and SA (pNITaraKm)] along with a recipient strain (KT2440Gm) were cultured in liquid medium, subsequently washed, mixed, and spotted onto solid 1/3 LB agar plates with various concentrations of arabinose. Following 24 hours of incubation, cells were collected and spread on selective plates containing tetracycline and gentamicin to count transconjugants. The displayed values represent the mean of two independent experiments, with error bars indicating the standard deviation.

### Transfer of ICE_KKS102_Tn*4677* is activated in *bphS* mutant

Conjugal transfer of ICE_KKS102_Tn*4677* in wildtype strain KKS102 has been reported to occur at a very low rate even in SA10 (13). As we have established the activating function of *traR* and *traR* upregulation in SA10, we reevaluated the transfer frequency of ICE using SA4 and SA10. In our experiment using KT2440Gm as a recipient, we observed no transconjugant when SA4 was used as a donor (transfer frequency <4.2×10^-9^). In contrast, when SA10 was used as a donor, transconjugants were obtained at a frequency of 3.8±3.6×10^-8^/input donor cell (Table 2).

**Table 2.** Transfer frequency of ICE_KKS102_Tn*4677* in SA4 and SA10.

| donor | recipient | selection | frequency / input donor cell |
| --- | --- | --- | --- |
| SA4 | KT2440Gm | Tc<sup>r</sup>, Gm<sup>r</sup> | $<4.2 \pm 1.4 \times 10^{-9}$ (n = 5) |
| SA10 | KT2440Gm | Tc<sup>r</sup>, Gm<sup>r</sup> | $3.8 \pm 3.6 \times 10^{-8}$ (n = 5) |

### TraR overexpression activated retransfer of ICE_KKS102_Tn*4677* in KT2440

We next examined whether transfer of ICE_KKS102_Tn*4677* in a KT2440-derived transconjugant, KT2440ICE, is activated by *traR* overexpression. KT2440ICE is a KT2440 derived transconjugant, and sensitive to gentamycin, which was selected for chloramphenicol resistance (an intrinsic phenotype of *P. putida* KT2440 (24)) and tetracycline resistance. As expected, mating of KT2440ICE (pNITaraKm_traR) with KT2440Gm resulted in the transfer of ICE. No transfer was observed in the vector control experiment using KT2440ICE (pNITaraKm). Transconjugants were obtained only when the experiment was conducted in the presence of arabinose (Table 3). These results indicated that TraR target is located on ICE_KKS102_Tn*4677*.

**Table 3.**
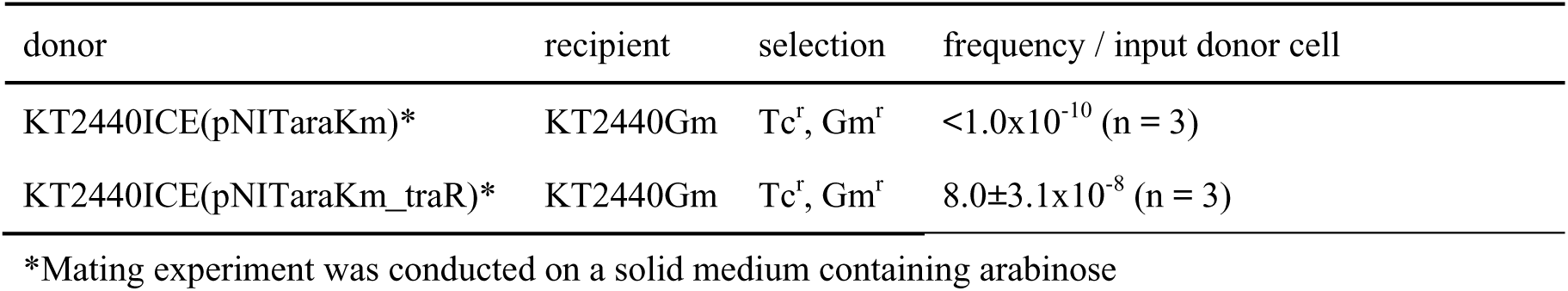
Transfer frequency of ICE_KKS102_Tn*4677* in KT2440ICE.

### Excision of ICE by *traR* overexpression

The increased transfer indicated enhanced ICE excision. The effect of *traR* overexpression on the *attP* and *attB* levels, which are formed by ICE excision, was investigated (Fig. 4). To note, in our preparative experiment using SA4 (pNITaraKm_traR) grown in liquid media, the increase in the level of excision was very modest and remained less than 2-fold. In search of the conditions, we found that we can observe marked increase when we spot and grow the cells on a solid medium. The conditions are very similar to those we employed in the mating experiments. Even on solid media, both *attP* and *attB* levels remained relatively low for the initial 12 hours of incubation after induction of *traR* expression by spotting the cells on arabinose-containing medium. Thereafter, *attP* and *attB* levels increased much more considerably reaching 130 and 40 folds increase for *attP* (panel B) and *attB* (panel C), respectively.

**Fig. 4:**
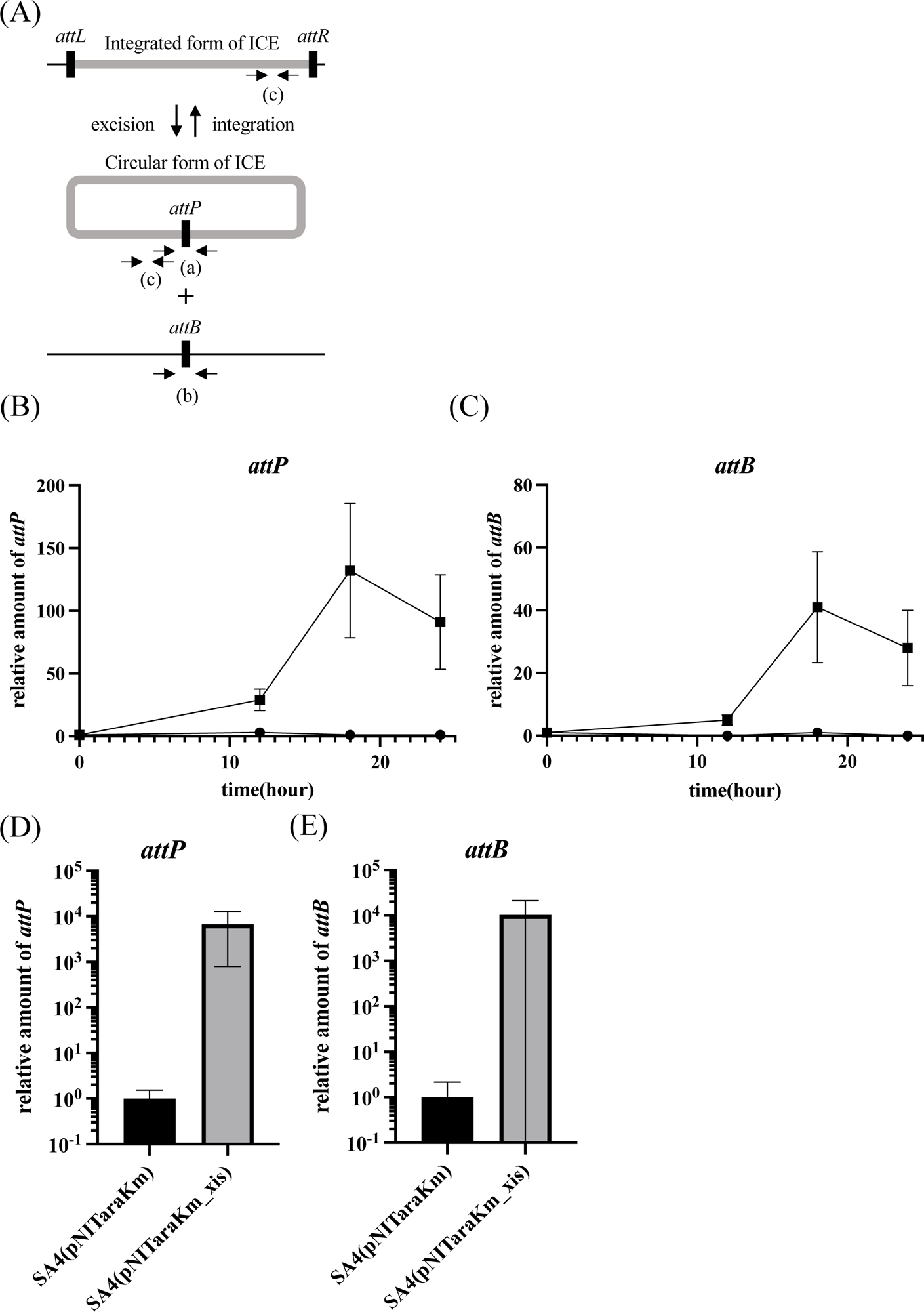
Overexpression of *traR* increased excised form of ICE. (A) Schematic representation of the circular and integrated forms of ICE. Three primer sets were designed to detect the circular form (*attP*, primer set a), a site after excision of ICE (*attB*, primer set b), and both forms (primer set c). (B, C) ICE excision by *traR* indcution. Strains SA4 (pNITaraKm) (solid circle) and SA4 (pNITaraKm_traR) (solid square) were cultured in liquid medium, harvested, and spotted onto solid 1/3 LB agar plates containing 10 mM arabinose. At each time point, cells were collected and subjected to qPCR analysis to quantify the amounts of *attP* (B) and *attB* (C) DNAs. The quantity of DNA was normalized using primers that target both forms of ICE. The values are relative value with the initial time point value set as 1. (D, E) ICE excision by *xis* indcution. Strains SA4 (pNITaraKm) and SA4 (pNITaraKm_xis) were cultured in liquid medium, harvested, and spotted onto solid 1/3 LB agar plates containing 10 mM arabinose. After one hour, cells were collected to quantify the amounts of *attP* (D) and *attB* (E) DNAs.

### *traR* overexpression resulted in up regulation of *xis*

We investigated whether the transcription levels of several genes of ICE, including the *xis* gene, are upregulated by the overexpression of *traR*. We found that increase in *xis* transcript level was observed only after prolonged incubation in the presence of arabinose (see discussion for details), which is consistent with the excision frequency data shown above. Fig. 5 shows the results of qRT-PCR, conducted using cells after 18 hours of incubation on a solid medium. Among the genes tested (*xis*, *int*, *traG*, and *traI*), the transcription level of *xis* increased 80-fold by inducing *traR*. The transcription levels of the *traG* and *int* increased 3-folds and 2-folds, respectively, and *traI* level remained unchanged.

**Fig. 5:**
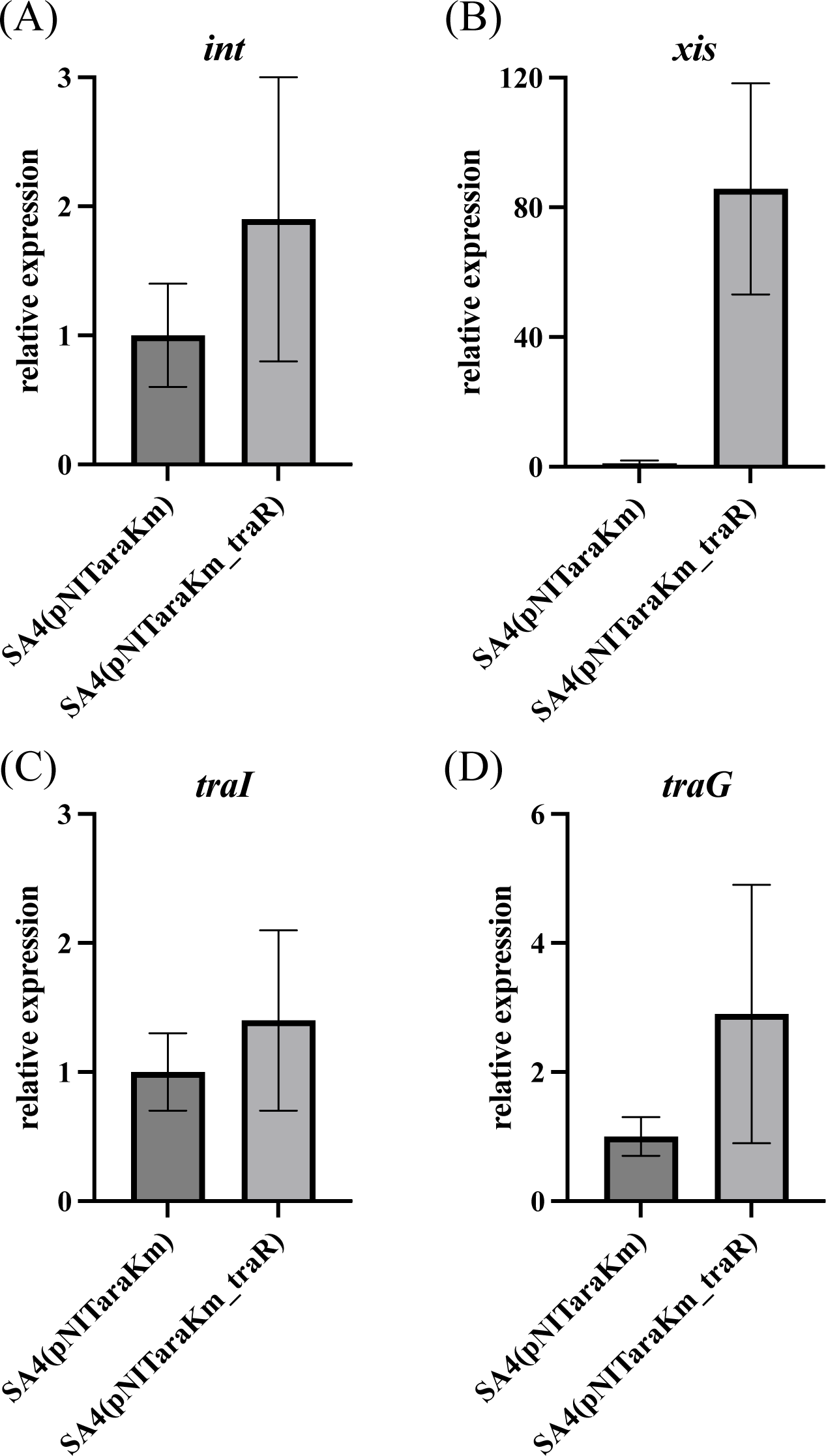
Transcriptional induction of ICE gene by overexpression of *traR*. Two strains SA4 (pNITaraKm) and SA4 (pNITaraKm_traR) were cultured in liquid medium, harvested, and spotted onto solid 1/3 LB agar plates containing arabinose. After 18 hours, cells were collected for RNA isolation, and qRT-PCR experiments were conducted for *int* (A), *xis* (B), *traI* (C), and *traG* (D). The values are expressed as relative to the strain SA4(pNITaraKm), with the *rrn* gene used as the internal standard. The values represent the average of three independent experiments, and error bars indicate standard deviations.

### Stimulation of excision by *xis* overexpression is not sufficient to activate transfer

To test if TraR mediated ICE excision, by upregulation of *xis*, is sufficient for the activation of ICE transfer, mating assay was conducted using strain SA4 carrying *xis* expression plasmid pNITaraKm_xis. As shown in Fig. 4, overexpression of *xis* resulted in the enhanced excision. However, no ICE transfer was observed (Table 4 panels D and E), indicating that TraR has more essential function than upregulation of *xis* to increase circular form ICE.

**Table 4.** Transfer frequency of pICEoriT.

| donor | recipient | selection | frequency / input donor cell |
| --- | --- | --- | --- |
| SA4 (pNITaraKm_xis) | KT2440Gm | Tc<sup>r</sup>, Gm<sup>r</sup> | $<3.6 \pm 0.92 \times 10^{-9}$ (n = 3) |
| SA4 (pNITaraKm) | KT2440Gm | Tc<sup>r</sup>, Gm<sup>r</sup> | $<5.0 \times 10^{-9}$ (n = 3) |
| SA4 (pNITaraKm_TraR) | KT2440Gm | Tc<sup>r</sup>, Gm<sup>r</sup> | $3.5 \pm 4.5 \times 10^{-7}$ (n=3) |

\*Mating experiment was conducted on a solid medium containing arabinose

### in silco search for TraR binding sites

We explored the DNA sequence upstream of *xis* to identify the binding sites of TraR. As shown in Fig. 6, at the immediate upstream region of -35 box element of a predicted promoter we found an interrupted inverted repeat (<u>CGTTT**T**G</u>ATGGAGACG<u>C**A**AAACG,</u> designated as BS^xis^) containing the LysR binding motif of TN_11_A (25). The distance of BS^xis^ from the -35 box of the promoter suggested that this is a canonical promoter regulated by a LysR-type transcriptional regulator. In searching for similar sequences, we identified a similar putative binding site (TT<u>T</u>A<u>T**T**G</u>AATCATTGG<u>C**A**AAACG</u>, designated as BS^15836^) upstream of a gene encoding hypothetical protein (C380_15836). BS^15836^ was also found immediate upstream of a predicted promoter. Further extensive search for similar sequences revealed a half binding site at upstream of *traG* putative promoter (CAAAACG designated as BS^traG^). Additionally, we found CAAAAGG (designated as BS^oriT^) within the well-conserved 20 bp sequence of *oriT* in ICE_KKS102_Tn*4677*, situated upstream of *traI* (see below). Also putative *traR* promoter was preceded by less conserved half site of CAAAA (designated as BS^traR^).

**Fig. 6.**
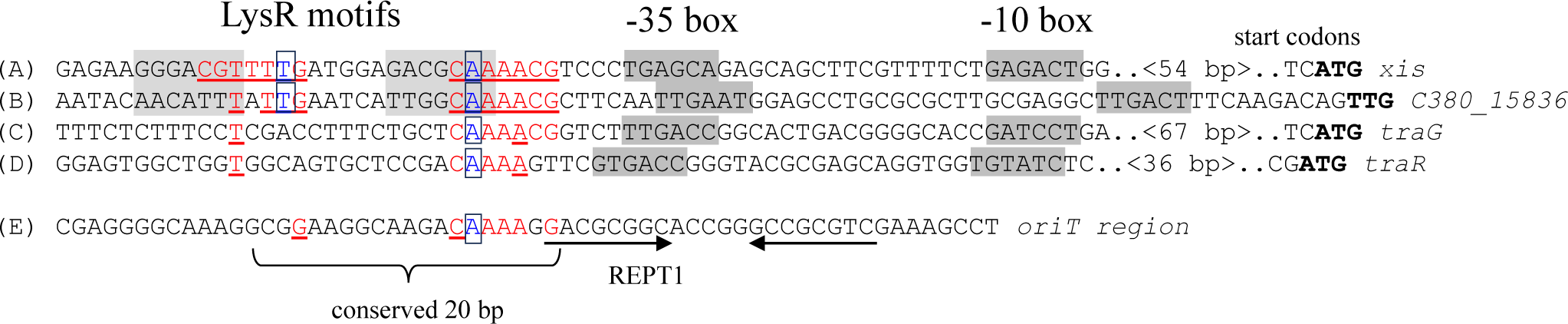
DNA sequences harboring putative TraR binding sites. A TN11A LysR motif, featuring T and A in an interrupted inverted repeat, was found immediately upstream of the predicted *xis* promoter (A). A similar sequence, with 3 mismatches across 14 nucleotides (matching nucleotides are underlined), was identified immediately upstream of the predicted promoter for C380-15836 (B). The half-site was located immediately upstream of the predicted promoters for *traG* (C) and *traR* (D). This half-site also appears in the 20-bp sequence conserved in the *traF*-*traI* intergenic region across related ICE elements (E). The start codon for each gene is highlighted in bold. Promoter elements (the -10 and -35 boxes), as predicted by the method described by Mulligan (38), are displayed on a shaded background. T and A in the LysR motif (TN_11_A) are enclosed in boxes. The REPT1 sequence is also illustrated.

### pICEoriT transferred to KT2440 by *traR* overexpression

To better understand the ICE transfer, it is essential to identify *oriT* of ICE_KKS102_Tn*4677*. Due to the high level of sequence conservation upstream of *traI* and downstream of *traF*, we predict that the *oriT* of ICE_KKS102_Tn*4677* is located in this intergenic region. To test this, a plasmid pICEoriT was constructed by cloning the intergenic region (463 bp) into a broad host range vector, pBBR1-MCS5 (26). The pICEoriT plasmid transferred to KT2440 not from SA7 or SA17 but from SA4 and SA10, with two orders of magnitudes higher for SA10 (Table 5), showing that *traR* activates the transfer of pICEoriT, and that *traR* is essential for transfer. Furthermore, overexpression of *traR* expressed from pNITaraKm_traR resulted in a frequency value of 1.9×10-3. The *traR* activation of transfer of pICEoriT, a circular plasmid entity, showed that *xis*-induction leading to ICE excision is not the sole role of TraR in transfer activation.

**Table 5.** Transfer frequency of pICEoriT.

| donor | recipient | selection | frequency / input donor cell |
| --- | --- | --- | --- |
| SA4 (pICEoriT) | KT2440 | Cm<sup>r</sup>, Gm<sup>r</sup> | $2.7 \pm 4.2 \times 10^{-7}$ (n = 2) |
| SA7 (pICEoriT) | KT2440 | Cm<sup>r</sup>, Gm<sup>r</sup> | $<2.9 \times 10^{-8}$ (n = 2) |
| SA10 (pICEoriT) | KT2440 | Cm<sup>r</sup>, Gm<sup>r</sup> | $1.1 \pm 0.7 \times 10^{-5}$ (n = 2) |
| SA17 (pICEoriT) | KT2440 | Cm<sup>r</sup>, Gm<sup>r</sup> | $<7.8 \times 10^{-8}$ (n = 2) |
| SA4 (pBBR1-MCS5) | KT2440 | Cm<sup>r</sup>, Gm<sup>r</sup> | $<1.1 \times 10^{-9}$ (n = 2) |

### Identification of essential *oriT* sequence

In the *oriT* region we found two LysR motifs (LysR motifs 1 and 2), conserved 20 bp sequence (see supplementary Fig. S1), and two repetitive elements (REPT1 and REPT2)(Fig. 7, panel A). To identify the minimum *oriT* region, we constructed a series of deletion plasmid of pICEoriT, and measured the transfer frequency under *traR*-overexpressing conditions. As shown in panel B in Fig. 7, transfer frequency of pICEoriT (3.1×10^-3^) was roughly unchanged when the *oriT* region was deleted from left side (*traF* side) up to position 261 (see del4 in Fig. 7), showing that the two LysR motifs are not important for *oriT* function. The frequency was decreased to undetectable level when deleted to position 298 (del8). When deleted from right side (*traI* side), deletion to position 393 (del6) did not affect the frequency, and when the REPT2 was deleted, the frequency decreased by 2 order of magnitude (del7). The deletion to just before the REPT1 did not abolish transfer (del9), but deletion of REPT1 resulted in undetectable level of frequency (del17). The del25 (carrying region from position 261 to 318) with the conserved 20 bp sequence and REPT1 maintained *oriT* activity. REPT1 and REPT2 sequences are not essential because substitution of bases (del9* and del24*) had some effects but did not abolish the *oriT* function. Altogether, we concluded that essential *oriT* functional sequence lies between position 261 to 318 (58 bp) where conserved 20 bp sequence and REPT1 are located, and that regions around RPET2 have some effect in enhancing transfer frequency.

**Fig. 7.**
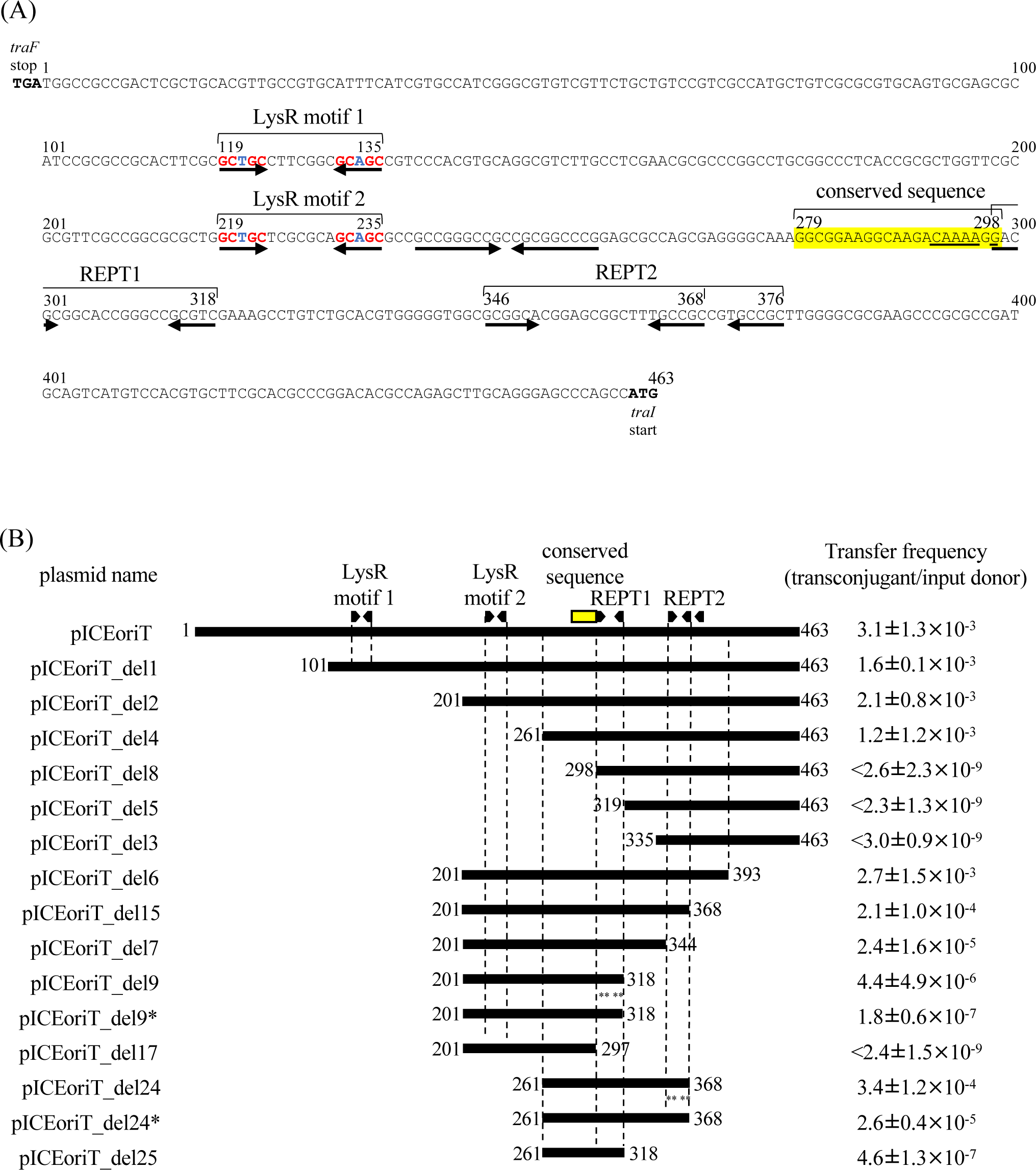
Deletion analysis of *traF*-*traI* intergenic region for *oriT* function. (A) DNA sequence of *traF*-*traI* intergenic region. The conserved 20 bp sequence, putative LysR motif, and repeat elements are indicated. The conserved 20-bp sequence is displayed on yellow background. (B) Deletion analysis of *oriT* region. The horizontal bars indicate DNA regions cloned into pBBR1-MCS. *oriT* plasmid names and transfer frequencies are show left and right to the bars, respectively.

## Discussion

In this study, we established that *traR,* co-transcribed with the *bph* operon, is the activator of ICE_KKS102_Tn*4677* transfer and identified *oriT* sequence. We also elucidated aspects of the mechanisms behind *traR*-mediated ICE transfer activation, in which TraR activates *xis* transcription leading to ICE excision. Also, upregulation of *traG* promoter by TraR was demonstrated. Our study shed light on the molecular and genetic basis of the dissemination of Tn*4371* family of ICEs (Fig. 8).

**Fig. 8.**
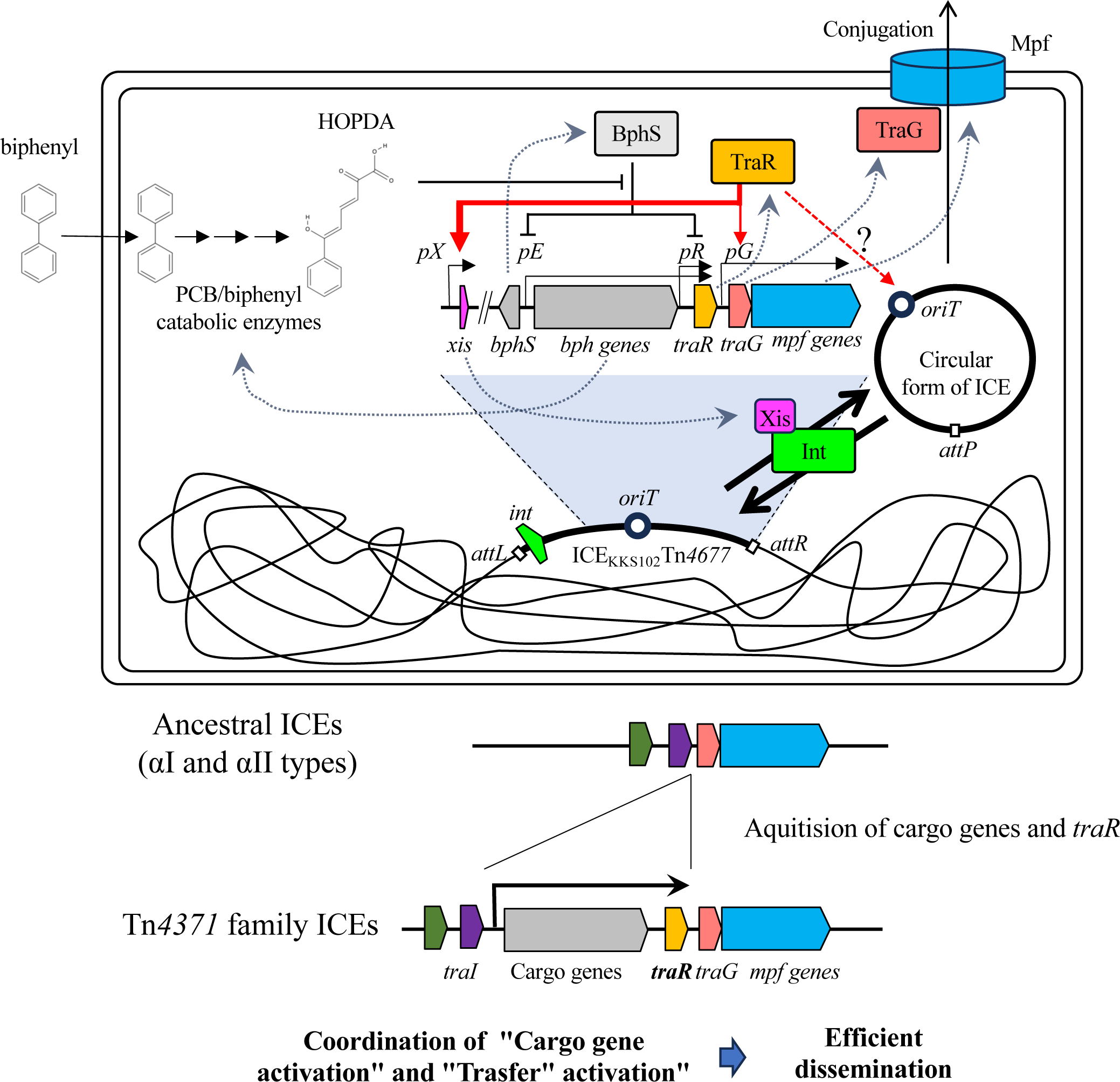
A model illustrating the activation of ICE transfer in response to biphenyl. In the presence of PCBs/biphenyl, the intermediate compound HOPDA mitigates repression by BphS, leading to the upregulation of the *pE* and *pR* promoters, which in turn results in the expression of TraR. TraR enhances the expression of *xis*, facilitating ICE excision. Additionally, TraR upregulates the *pG* promoter, inducing the expression of TraG (the coupling protein) and the formation of the mating pair. It is speculated that TraR may interact with *oriT*. Consequently, ICE transfer is activated in the presence of PCBs/biphenyl, in which TraR plays the central role. The *traR* gene direction and location situated downstream of cargo genes are conserved characteristics among Tn*4371* family ICEs. This gene organization allows the ICE transfer under environmental conditions that activate cargo gene transcription. This would increase the likelihood of the ICE amplifying its copy number, as the cargo genes are likely to confer fitness advantages to the host bacteria, thereby facilitating the efficient dissemination. Notably, ancestral ICEs, namely types αI and αII, do not possess the cargo and *traR* genes at the corresponding locations.

### *traR* transcription initiated from *pE* and *pR* promoters

*traR* is transcribed from two promoters, *pE* and *pR*. The *pE* promoter is regulated by BphS (16), and in this study we demonstrated an increase in *traR* transcript in the absence of *bphS*. In addition replacing *pE* promoter with a constitutive strong promoter (*tac* promoter (22)) resulted in increased *traR* transcript. These findings clearly showed that *traR* is transcribed from *pE* under control of BphS, which allows the *bph* operon for PCB/biphenyl degradation to be upregulated only in the presence of the degradation substrate. We also found that *pR* promoter is upregulated in the absence of *bphS*. Because a number of LysR-type transcriptional regulators are known to control its own transcription(25), we speculated that high *pR* activity in KLZ60ΔS is due to *traR* upregulation. To test this, KLZ60 was transformed with *traR* expression plasmid pNITaraKm_traR, which carries *traR* gene under arabinose-inducible pBAD promoter, and tested for LacZ activity. The effect of *traR* was tested by using cells grown on a solid medium, however we were unable to obtain any result showing that pR is upregulated by TraR.

### Implications of ICE-transfer activation by *bph* operon substrate

The regulatory system, which links the availability of the *bph* degradation system’s substrate to the active horizontal transfer of ICE, suggests a strategic advantage in the dissemination of ICE_KKS102_Tn*4677*. This system enhances the transfer of cargo *bph* genes to surrounding bacteria in environments where these genes confer a fitness advantage, potentially playing a crucial role in ICE dissemination. Such a mechanism ensures effective spread of the ICE, as bacterial cells in these environments are more likely to benefit from the acquired capabilities.

A wealth of studies has highlighted diverse mechanisms controlling bacterial horizontal gene transfer (HGT). As for the self-transmissible chlorocatechol degradative *clc* genes of *Pseudomonas knackmussii* strain B13, the clc element’s integrase gene expression was significantly stimulated by growing cultures on 3-chlorobenzoate (27). In *Bacillus subtilis*, the conjugation of a plasmid, pLS20, is inhibited at high levels of the quorum signaling peptide, which can be beneficial in preventing unnecessary gene transfer in densely populated environments (28). In a study by Beaber JW, bacterial SOS response induced by DNA-damage was shown to increase the frequency of transfer of SXT element, an ICE derived from *Vibrio cholerae*. (29)

Our finding that *traR*, whose transcription is co-upregulated with the conditional activation of cargo *bph* genes, activates the horizontal transfer of ICE_KKS102_Tn*4677*, is another good example of how the horizontal transfer of mobile genetic elements is regulated.

### Delay of *xis* upregulation after *traR* induction

We observed a considerable delay in the increase of *xis* transcription and the excision of ICE following the induction of *traR* with arabinose. In bacterial transcription induction, responses typically occur within a very short time span. For instance, with the pBAD promoter, transcription is known to increase within a short period after arabinose addition (30). However, when we induced *traR* expression with arabinose, we did not observe an increase in *xis* transcription within the usual time scale. In one RNAseq analysis, *traR* induction resulted in 122-folds increase of NGS reads mapped to *traR* gene one hour after addition of arabinose, yet no *xis* upregulation was observed.

Several factors might contribute to this delayed induction, although the precise reasons remain unclear. One possibility is that the stability of TraR or translation efficiency of *traR* is regulated during different growth phases. Upon examining TraR expression levels through Western blot analysis, we found that a significant amount of time was required before the TraR bands became observable.

Alternatively, it is worth noting that TraR is a LysR-type transcription factor, which might require specific effector for transcription activation. The unknown effector might be generated at very late stage of growth. Currently, the cues for identifying the unknown effector are quite limited. However, the fact that *traR*-mediated ICE excision did not occur in liquid culture may provide some hints. The unknown signal may be related to the stable contact with other cells through formation of T4SS, which might readily break from the cell surface (14). Further investigation is needed to fully understand the delay in TraR-mediated *xis* activation.

### DNA binding of TraR

To investigate the binding of TraR to the DNA regions upstream of *xis*, we purified an N-terminal His-tagged TraR protein and tested its binding to BSxis. Despite considerable efforts, only a very weak interaction was observed between TraR and the DNA fragment containing BSxis. The reason for this weak interaction remains unclear, but it may be attributed to either the stability of the TraR protein or the absence of an unknown effector. The addition of an N-terminal His-tag is unlikely to be the cause, as the N-terminal His-tagged TraR demonstrated functionality in increasing the ICE circular form. TraR binding to other putative binding sites were also not successful.

### Mechanisms of ICE transfer activation by TraR

In this study we showed that TraR activated *xis* transcription and resulted in enhanced ICE excision, however, excision alone was not sufficient for the ICE transfer to occur. The plasmid cloned with *oriT* (pICEoriT) required *traR* overexpression for transfer. In addition, while *xis* overexpression did enhance ICE’s circular form, it did not result in ICE transfer. The transfer activation of ICE_KKS102_Tn*467*7 in KT2440 by *traR* overexpression indicates that other TraR targets are located on the ICE. Although TraR overexpression induced a modest threefold increase in *traG* expression (Fig. 5), this upregulation suggests it’s crucial role in the activation of transfer. The presence of putative TraR half binding site very close to the *traG* promoter (*pG* promoter) suggests that the *pG* is under the control of TraR. In our preparative RNAseq analysis (n=1) exhibited that very shortly after *traR* induction (one hour), transcript of *traG* and *mpf* genes increased 1.9 folds. However, in the subsequent attempt, these genes were not upregulated after one hour possibly due to slight unintended differences in experimental conditions, suggesting that unknown-inducing signal, which was present in the initial attempt, is required to upregulate *pG*. Identifying the yet-unidentified TraR effector would lead to a better understanding of TraR-mediated *pG* activation, as well as overall mechanism of TraR mediated ICE transfer activation.

### Possible role of TraR as an accessory protein

Other TraR’s possible target is *oriT*. Although an *oriT* is a cis-acting DNA region required for the conjugation of plasmids and ICEs, it had not been identified in the ICEs of the Tn*4371* family. In this study, we experimentally demonstrated that the intergenic region between the *traI* and *traF* genes on ICE_KKS102_Tn*4766* functions as an *oriT*. This region was not detected by oriTfinder (31), a software tool designed for predicting *oriTs* from DNA sequence information, indicating that the *oriT* is distinct from previously identified *oriTs* and represents a novel *oriT*.

The *nic* site is most likely located within the 20-bp yellow-highlighted conserved region, because this area is required for DNA transfer (Fig. 7), and *nic* sites typically occur around 10 nucleotides away from an inverted repeat (32, 33). Within this 20 bp sequence, we found a sequence of CAAAAGG, a half site of the possible TraR binding sequence found upstream of *xis*. For some LysR-type transcriptional regulators, including CbnR, CysB, AlsR, and OccR, bending of DNA upon binding to their binding site are reported (34) (35) (36) (37). Further in vivo and in vitro studies are required to confirm whether TraR binds to this sequence and potentially bends DNA around *oriT*, acting as an accessory protein to aid TraI’s nicking function at this site.

### *int* gene upregulation might be the result of circular ICE formation

The modest increase in *int* gene transcription level (2 folds) might result from overexpression of *xis*, and subsequent formation of circular ICE. As described previously, *int* gene is highly transcribed in the circular form (13). A strong promoter located close to the right boundary of ICE transcribe *int* gene upon forming the circular form. Therefore, if a minor fraction of cells undergoes ICE excision due to *traR* overexpression, it’s possible for the overall *int* transcript levels to increase to this extent.

### Conservation of *traR* gene among Tn*4371* family ICEs

In Tn*4371* family ICEs, predominantly found in β and γ proteobacteria, the *traR* gene is situated between the accessory region 3 and *mpf* genes, oriented in the same direction as the *mpf* genes. This gene organization is not seen in the related ICEs from α proteobacteria or plasmids, which were collected based on similarity of encoded TraG to TraG from KKS102 (13).

### Role of TraR in the emergence of Tn*4371* family ICEs

A phylogenic analysis of Tn*4371* family ICEs and related ICEs outside of Tn4371 family (types αI and αII) using TraG amino acid sequences showed that branch lengths among Tn*4371* family ICEs are shorter than those among the related elements from α proteobacteria or plasmids (13), suggesting rapid dissemination of Tn*4371* family ICEs. In addition, it seemed that Tn*4371* family ICEs have emerged from ancestral ICEs with gene organization of types αI and αII (13).

In most of Tn*4371* family ICEs, direction of genes in the accessory region 3 are heading toward the *traR* (Fig. 8), i.e., cargo gene transcription is coordinated with *traR* transcription. Supplementary Fig.8 shows that 30 out of 32 ICEs founding the Tn*4371* family (13) exhibit this organization. On the other hand, *traR* is absent from this location in Tn*4371*-related ICEs of types αI and αII. From an evolutionary perspective, with this organization of Tn*4371* family ICEs, incorporation of a new cargo gene into accessory region 3 readily setup this coordinated transcription. This coordination would be advantageous for the ICE dissemination because the environmental conditions where cargo genes are activated are likely the conditions where the ICE gives the fitness advantage to the host bacteria thereby giving more chance to increase ICE’s copy number. Although how ICEs acquire new cargo genes remains totally unknown, we propose that the recruitment of *traR* gene to the current position to the ancestral ICE have established ICE Tn*4371* family by enabling efficient dissemination due to the coordinated transcription of cargo genes and *traR* gene.

## DATA AVAILABILITY

The original data supporting the findings of this study are available upon reasonable request. To request the data, please contact Yoshiyuki Ohtsubo at. The data will be made available under the condition that it is used only for non-commercial research purposes and that any publications or presentations resulting from the use of the data cite this study as the source of the data.

## Acknowledgment

This work was supported by JST, the establishment of university fellowships towards the creation of science technology innovation, Grant Number JPMJFS2102.

## Author contributions

S.M. performed most of the experiments and drafted initial manuscript. K.S. performed the initial *traR* overexpression experiment. S.N. participated in conducting in designing and conducting part of experiments. Y.O. supervised the whole project and completed the manuscript. All authors, S.M., K.K., S.N., K.S., M.T., Y.N., and Y.O. contributed to discussions, editing, and revision of the manuscript.

## REFERENCES

1. Roberts AP, Mullany P. 2011. Tn916-like genetic elements: a diverse group of modular mobile elements conferring antibiotic resistance. FEMS Microbiol Rev 35:856–71.

2. Partridge SR, Kwong SM, Firth N, Jensen SO. 2018. Mobile Genetic Elements Associated with Antimicrobial Resistance. Clinical Microbiology Reviews 31.

3. Durrant MG, Li MM, Siranosian BA, Montgomery SB, Bhatt AS. 2020. A Bioinformatic Analysis of Integrative Mobile Genetic Elements Highlights Their Role in Bacterial Adaptation. Cell Host Microbe 27:140–153 e9.

4. Top EM. 2002. Transfer of Catabolic Plasmids in Soil and Activated Sludge: A Feasible Bioaugmentation Strategy?, p 91–103. In Agathos SN, Reineke W (ed), Biotechnology for the Environment: Strategy and Fundamentals doi:10.1007/978-94-010-0357-5_6. Springer Netherlands, Dordrecht.

5. Wozniak RA, Waldor MK. 2010. Integrative and conjugative elements: mosaic mobile genetic elements enabling dynamic lateral gene flow. Nat Rev Microbiol 8:552–63.

6. Bragagnolo N, Rodriguez C, Samari-Kermani N, Fours A, Korouzhdehi M, Lysenko R, Audette GF. 2020. Protein Dynamics in F-like Bacterial Conjugation. Biomedicines 8.

7. Springael D, Kreps S, Mergeay M. 1993. Identification of a catabolic transposon, Tn4371, carrying biphenyl and 4-chlorobiphenyl degradation genes in Alcaligenes eutrophus A5. J Bacteriol 175:1674–81.

8. Toussaint A, Merlin C, Monchy S, Benotmane MA, Leplae R, Mergeay M, Springael D. 2003. The biphenyl- and 4-chlorobiphenyl-catabolic transposon Tn4371, a member of a new family of genomic islands related to IncP and Ti plasmids. Appl Environ Microbiol 69:4837–45.

9. Ryan MP, Pembroke JT, Adley CC. 2009. Novel Tn*4371*-ICE like element in *Ralstonia pickettii* and genome mining for comparative elements. BMC Microbiol 9:242.

10. Hirose J, Fujihara H, Watanabe T, Kimura N, Suenaga H, Futagami T, Goto M, Suyama A, Furukawa K. 2019. Biphenyl/PCB Degrading bph Genes of Ten Bacterial Strains Isolated from Biphenyl-Contaminated Soil in Kitakyushu, Japan: Comparative and Dynamic Features as Integrative Conjugative Elements (ICEs). Genes (Basel) 10.

11. Qian C, Liu H, Cao J, Ji Y, Lu W, Lu J, Li A, Zhu X, Shen K, Xu H, Chen Q, Zhou W, Lu H, Lin H, Zhang X, Li Q, Lin X, Li K, Xu T, Zhu M, Bao Q, Zhang H. 2021. Identification of floR Variants Associated With a Novel Tn4371-Like Integrative and Conjugative Element in Clinical Pseudomonas aeruginosa Isolates. Front Cell Infect Microbiol 11:685068.

12. Hirose J. 2023. Diversity and Evolution of Integrative and Conjugative Elements Involved in Bacterial Aromatic Compound Degradation and Their Utility in Environmental Remediation. Microorganisms 11.

13. Ohtsubo Y, Ishibashi Y, Naganawa H, Hirokawa S, Atobe S, Nagata Y, Tsuda M. 2012. Conjugal Transfer of Polychlorinated Biphenyl/Biphenyl Degradation Genes in *Acidovorax* sp. Strain KKS102, Which Are Located on an Integrative and Conjugative Element. J Bacteriol 194:4237–48.

14. Costa TRD, Patkowski JB, Macé K, Christie PJ, Waksman G. 2023. Structural and functional diversity of type IV secretion systems. Nature Reviews Microbiology doi:10.1038/s41579-023-00974-3.

15. Ohtsubo Y, Maruyama F, Mitsui H, Nagata Y, Tsuda M. 2012. Complete genome sequence of *Acidovorax* sp. strain KKS102, a polychlorinated-biphenyl degrader. J Bacteriol 194:6970–1.

16. Ohtsubo Y, Delawary M, Kimbara K, Takagi M, Ohta A, Nagata Y. 2001. BphS, a key transcriptional regulator of *bph* genes involved in polychlorinated biphenyl/biphenyl degradation in *Pseudomonas* sp. KKS102. J Biol Chem 276:36146–54.

17. Ohtsubo Y, Nagata Y, Kimbara K, Takagi M, Ohta A. 2000. Expression of the *bph* genes involved in biphenyl/PCB degradation in *Pseudomonas* sp. KKS102 induced by the biphenyl degradation intermediate, 2-hydroxy-6-oxo-6-phenylhexa-2,4-dienoic acid. Gene 256:223–8.

18. Ohtsubo Y, Goto H, Nagata Y, Kudo T, Tsuda M. 2006. Identification of a response regulator gene for catabolite control from a PCB-degrading beta-proteobacteria, *Acidovorax* sp. KKS102. Mol Microbiol 60:1563–75.

19. Sakai K, Kishida K, Matsumoto S, Nagata Y, Tsuda M, Ohtsubo Y. 2024. Three distinct metabolic phases of PCB/biphenyl degrader Acidovorax sp. KKS102 in nutrient broth. Biosci Biotechnol Biochem doi:10.1093/bbb/zbad178.

20. Sato T, Nonoyama S, Kimura A, Nagata Y, Ohtsubo Y, Tsuda M. 2017. The Small Protein HemP Is a Transcriptional Activator for the Hemin Uptake Operon in Burkholderia multivorans ATCC 17616. Appl Environ Microbiol 83.

21. Ono A. 2007. 培養非依存的手法による土壌からの環境汚染物質分解酵素遺伝子の取得と解析[Acquisition and Analysis of Environmental Pollutant-Degrading Enzyme Genes from Soil via Culture-Independent Methods].p129-p130.

22. Amann E Fau - Brosius J, Brosius J Fau - Ptashne M, Ptashne M. Vectors bearing a hybrid trp-lac promoter useful for regulated expression of cloned genes in Escherichia coli.

23. Ohtsubo Y, Shimura M, Delawary M, Kimbara K, Takagi M, Kudo T, Ohta A, Nagata Y. 2003. Novel approach to the improvement of biphenyl and polychlorinated biphenyl degradation activity: promoter implantation by homologous recombination. Appl Environ Microbiol 69:146–53.

24. Fernandez M, Conde S, de la Torre J, Molina-Santiago C, Ramos JL, Duque E. 2012. Mechanisms of resistance to chloramphenicol in Pseudomonas putida KT2440. Antimicrob Agents Chemother 56:1001–9.

25. Schell MA. 1993. Molecular biology of the LysR family of transcriptional regulators. Annu Rev Microbiol 47:597–626.

26. Kovach ME, Elzer PH, Hill DS, Robertson GT, Farris MA, Roop RM, 2nd, Peterson KM. 1995. Four new derivatives of the broad-host-range cloning vector pBBR1MCS, carrying different antibiotic-resistance cassettes. Gene 166:175–6.

27. Sentchilo V, Ravatn R, Werlen C, Zehnder AJ, van der Meer JR. 2003. Unusual integrase gene expression on the clc genomic island in Pseudomonas sp. strain B13. J Bacteriol 185:4530–8.

28. Singh PK, Serrano E, Ramachandran G, Miguel-Arribas A, Gago-Cordoba C, Val-Calvo J, Lopez-Perez A, Alfonso C, Wu LJ, Luque-Ortega JR, Meijer WJJ. 2020. Reversible regulation of conjugation of Bacillus subtilis plasmid pLS20 by the quorum sensing peptide responsive anti-repressor RappLS20. Nucleic Acids Res 48:10785–10801.

29. Beaber JW, Hochhut B, Waldor MK. 2004. SOS response promotes horizontal dissemination of antibiotic resistance genes. Nature 427:72–4.

30. Guzman LM, Belin D, Carson MJ, Beckwith J. 1995. Tight regulation, modulation, and high-level expression by vectors containing the arabinose PBAD promoter. J Bacteriol 177:4121–30.

31. Li X, Xie Y, Liu M, Tai C, Sun J, Deng Z, Ou HY. 2018. oriTfinder: a web-based tool for the identification of origin of transfers in DNA sequences of bacterial mobile genetic elements. Nucleic Acids Res 46:W229–W234.

32. Zrimec J, Lapanje A. 2018. DNA structure at the plasmid origin-of-transfer indicates its potential transfer range. Sci Rep 8:1820.

33. Kishida K, Inoue K, Ohtsubo Y, Nagata Y, Tsuda M. 2017. Host Range of the Conjugative Transfer System of IncP-9 Naphthalene-Catabolic Plasmid NAH7 and Characterization of Its oriT Region and Relaxase. Appl Environ Microbiol 83.

34. Giannopoulou EA, Senda M, Koentjoro MP, Adachi N, Ogawa N, Senda T. 2021. Crystal structure of the full-length LysR-type transcription regulator CbnR in complex with promoter DNA. FEBS J 288:4560–4575.

35. Hryniewicz MM, Kredich NM. 1994. Stoichiometry of binding of CysB to the cysJIH, cysK, and cysP promoter regions of Salmonella typhimurium. J Bacteriol 176:3673–82.

36. Akakura R, Winans SC. 2002. Constitutive mutations of the OccR regulatory protein affect DNA bending in response to metabolites released from plant tumors. J Biol Chem 277:5866–74.

37. Hartig E, Fradrich C, Behringer M, Hartmann A, Neumann-Schaal M, Jahn D. 2018. Functional definition of the two effector binding sites, the oligomerization and DNA binding domains of the Bacillus subtilis LysR-type transcriptional regulator AlsR. Mol Microbiol 109:845–864.

38. Mulligan ME, Hawley DK, Entriken R, McClure WR. 1984. Escherichia coli promoter sequences predict in vitro RNA polymerase selectivity. Nucleic Acids Res 12:789–800.

39. Katoh K, Rozewicki J, Yamada KD. 2019. MAFFT online service: multiple sequence alignment, interactive sequence choice and visualization. Brief Bioinform 20:1160–1166.

40. Ohtsubo Y, Ikeda-Ohtsubo W, Nagata Y, Tsuda M. 2008. GenomeMatcher: a graphical user interface for DNA sequence comparison. BMC Bioinformatics 9:376.

41. Kimbara K, Hashimoto T, Fukuda M, Koana T, Takagi M, Oishi M, Yano K. 1988. Isolation and characterization of a mixed culture that degrades polychlorinated biphenyls. Agric Biol Chem 52:2885–2891.

42. Nelson KE, Weinel C, Paulsen IT, Dodson RJ, Hilbert H, Martins dos Santos VA, Fouts DE, Gill SR, Pop M, Holmes M, Brinkac L, Beanan M, DeBoy RT, Daugherty S, Kolonay J, Madupu R, Nelson W, White O, Peterson J, Khouri H, Hance I, Chris Lee P, Holtzapple E, Scanlan D, Tran K, Moazzez A, Utterback T, Rizzo M, Lee K, Kosack D, Moestl D, Wedler H, Lauber J, Stjepandic D, Hoheisel J, Straetz M, Heim S, Kiewitz C, Eisen JA, Timmis KN, Dusterhoft A, Tummler B, Fraser CM. 2002. Complete genome sequence and comparative analysis of the metabolically versatile Pseudomonas putida KT2440. Environ Microbiol 4:799–808.

43. Miyazaki R, Ohtsubo Y, Nagata Y, Tsuda M. 2008. Characterization of the traD operon of naphthalene-catabolic plasmid NAH7: a host-range modifier in conjugative transfer. J Bacteriol 190:6281–9.

44. Heeb S, Itoh Y, Nishijyo T, Schnider U, Keel C, Wade J, Walsh U, O’Gara F, Haas D. 2000. Small, stable shuttle vectors based on the minimal pVS1 replicon for use in gram-negative, plant-associated bacteria. Mol Plant Microbe Interact 13:232–7.

45. Nagayama H, Sugawara T, Endo R, Ono A, Kato H, Ohtsubo Y, Nagata Y, Tsuda M. 2015. Isolation of oxygenase genes for indigo-forming activity from an artificially polluted soil metagenome by functional screening using Pseudomonas putida strains as hosts. Appl Microbiol Biotechnol 99:4453–70.

46. Datsenko KA, Wanner BL. 2000. One-step inactivation of chromosomal genes in Escherichia coli K-12 using PCR products. Proc Natl Acad Sci U S A 97:6640–5.

